# Fungi of the order Mucorales express a “sealing-only” tRNA ligase

**DOI:** 10.1101/2023.11.16.567474

**Authors:** Khondakar Sayef Ahammed, Ambro van Hoof

## Abstract

Some eukaryotic pre-tRNAs contain an intron that is removed by a dedicated set of enzymes. Intron-containing pre-tRNAs are cleaved by tRNA splicing endonuclease (TSEN), followed by ligation of the two exons and release of the intron. Fungi use a “heal and seal” pathway that requires three distinct catalytic domains of the tRNA ligase enzyme, Trl1. In contrast, humans use a “direct ligation” pathway carried out by RTCB, an enzyme completely unrelated to Trl1. Because of these mechanistic differences, Trl1 has been proposed as a promising drug target for fungal infections. To validate Trl1 as a broad-spectrum drug target, we show that fungi from three different phyla contain Trl1 orthologs with all three domains. This includes the major invasive human fungal pathogens, and these proteins each can functionally replace yeast Trl1. In contrast, species from the order Mucorales, including the pathogens *Rhizopus arrhizus* and *Mucor circinelloides*, contain an atypical Trl1 that contains the sealing domain, but lack both healing domains. Although these species contain fewer tRNA introns than other pathogenic fungi, they still require splicing to decode three of the 61 sense codons. These sealing-only Trl1 orthologs can functionally complement defects in the corresponding domain of yeast Trl1 and use a conserved catalytic lysine residue. We conclude that Mucorales use a sealing-only enzyme together with unidentified non-orthologous healing enzymes for their heal and seal pathway. This implies that drugs that target the sealing activity are more likely to be broader-spectrum antifungals than drugs that target the healing domains.

## Introduction

In all eukaryotes, a subset of tRNA precursors contains introns that are removed by a specialized set of tRNA splicing enzymes (Popow et al. 2012; Phizicky and Hopper 2023). In contrast to spliceosomal splicing of mRNAs and other non-coding RNAs, tRNA splicing is a protein-mediated process initiated by cleavage of the pre-tRNAs by the tRNA splicing endonuclease complex (TSEN) (Peebles et al. 1979; Hayne et al. 2023). TSEN generates a 5’ exon that ends with a 2’ 3’ cyclic phosphate and a 3’ exon with a 5’ hydroxyl in a catalytic mechanism that is similar to that used by RNase A (Peebles et al. 1983). This initial step of tRNA splicing appears conserved in all eukaryotes, but two distinct mechanisms are used for the subsequent ligation step of two tRNA halves (5’ and 3’ exons) (Greer et al. 1983; Phizicky et al. 1986).

Most Metazoa and some protozoa use a “direct ligation” mechanism to join the tRNA exons, which is carried out by RTCB enzymes. RCTB uses a catalytic histidine residue to transfer GMP onto the 3’ end of the 5’ exon, generating an activated exon that can then be ligated to the 3’ exon (Chakravarty et al. 2012; Englert et al. 2012; Desai et al. 2013; Jacewicz et al. 2022). RTCB has been implicated in tRNA splicing in humans, *Caenorhabditis elegans*, and *Drosophila melanogaster*, and homologs of RTCB are encoded in most metazoan genomes (Popow et al. 2011; Kosmaczewski et al. 2014; Lu et al. 2014; Nandy et al. 2017). Although bacteria do not have tRNA introns (except rare group I self-splicing introns), some bacteria, including *Escherichia coli* also contain RTCB homologs that instead function in tRNA repair (Tanaka et al. 2011; Tanaka and Shuman 2011; Chen and Wolin 2023).

In contrast, most fungi and plants and some other protozoa use a “heal and seal” mechanism carried out by the multifunctional enzyme Trl1 to join tRNA exons (Konarska et al. 1981; Greer et al. 1983; Phizicky et al. 1986; Abelson et al. 1998; Sawaya et al. 2003; Lopes et al. 2015; Phizicky and Hopper 2023). The heal and seal mechanism involves three distinct steps that are carried out by three different catalytic domains of Trl1. In one “healing” reaction, the cyclic phosphodiesterase (CPD) domain converts the 2’3’ cyclic phosphate of the 5’ exon to a 3’-OH, 2’ PO_4_ end. In a second “healing” reaction, the kinase domain converts the 5’ hydroxyl of the 3’ exon to a 5’ phosphate. Finally, in the “sealing” reaction, the ligase domain uses a catalytic lysine residue to transfer AMP onto the 3’ exon, which activates it for ligation to the 5’ exon (Wang et al. 2006; Banerjee et al. 2019). Thus, both the catalytic residue (His *vs.* Lys) that carries out activation, as well as the exon (5’ *vs.* 3’) that is activated, differ between the direct ligation and heal and seal pathways. The heal and seal mechanism generates an initial ligation product with a 2’ phosphate at the exon junction, which is removed by tRNA 2’-phosphotransferase (Tpt1) (Spinelli et al. 1997; Steiger et al. 2001). The bacteriophage T4 encodes a polynucleotide kinase (PNK) and RNA ligase (Rnl1) that are widely used in molecular biology but whose normal role is in a related but distinct tRNA repair pathway (Amitsur et al. 1987; Shuman 2023). The T4 pathway does not involve a 2’PO_4_ intermediate or a Tpt1. Strikingly, these distinct pathways are interchangeable and the yeast *TRL1* gene can be replaced by either *E. coli* RtcB or T4 PNK and T4 Rnl1 (Schwer et al. 2004; Tanaka et al. 2011).

Both RTCB and Trl1 have an additional function in the splicing of a single mRNA during unfolded protein response (UPR) (Sidrauski et al. 1996; Gonzalez et al. 1999; Mori 2009). During endoplasmic reticulum (ER) stress, the endonuclease Ire1 cleaves the *Saccharomyces cerevisiae* (hereafter “yeast”) *HAC1* and human *XBP1* mRNAs. Similar to TSEN, Ire1 also generates a 5’ exon that ends with a 2’ 3’ cyclic phosphate and a 3’ exon with a 5’ hydroxyl, and these exons are ligated by RTCB and Trl1, in Metazoa and Fungi, respectively, using the same mechanism used for tRNA ligation.

The fungal tRNA exon ligation mechanism has been proposed as an attractive target for anti-fungal drug development for several reasons (Wang and Shuman 2005; Tanaka et al. 2011; Chakravarty et al. 2012; Remus et al. 2016): (i) tRNA splicing is essential in all eukaryotes. Without it, eukaryotes cannot generate a complete set of tRNAs and cannot translate a subset of codons. (ii) Fungi are relatively closely related to humans and there are very few fundamental differences in their biochemical pathways. As a result, there are very few genes that are essential in fungi but that do not have a close homolog in the human genome (Braun et al. 2005; Liu et al. 2006). (iii) The biochemical mechanism of tRNA exon ligation by fungal Trl1 is distinct from the metazoan enzyme RTCB. (iv) All three domains of Trl1 are required for its function in tRNA ligation and cell survival in the model fungus *Saccharomyces cerevisiae* (Sawaya et al. 2003; Wang and Shuman 2005; Wang et al. 2006). Therefore, a Trl1 inhibitor that targets any of the three essential domains is a potential anti-fungal drug candidate.

Novel antifungal drugs are urgently needed. There are only three classes of approved antifungal drugs for invasive disease, and resistance to these drugs limits their use. The CDC considers *C. auris* as one of six urgent threats of antimicrobial resistant pathogens, other drug-resistant *Candida* species as a serious threat, and azole-resistant *Aspergillus* as a watch list threat (https://www.cdc.gov/drugresistance/biggest-threats.html). In the US, fungal disease are responsible for seven billion dollars in annual health care costs (Benedict et al. 2019) and 3% of all hospitalized patients receive an antifungal drug (Vallabhaneni et al. 2018). The WHO issued a report last year calling for increased research into 19 fungi that represent the greatest threat to human health (https://www.who.int/news/item/25-10-2022-who-releases-first-ever-list-of-health-threatening-fungi). Fungal infections alarmingly have become one of the major global health problems among immunocompromised individuals. Among several types of fungal infections, invasive and pulmonary fungal diseases such as candidiasis, aspergillosis, cryptococcal meningitis, pneumocystis pneumonia, histoplasmosis, and mucormycosis are leading causes of morbidity and mortality worldwide (Garber 2001; von Lilienfeld-Toal et al. 2019; Kainz et al. 2020; Reddy et al. 2022). In addition, patients with existing health conditions such as HIV, cancer, and SARS-CoV-2 appear to have a high risk of fungal co-infections (Limper (Bodey et al. 1992; Limper et al. 2017; Soltani et al. 2022). Strikingly, a sharp increase in deaths caused by fungal infection was observed during the coronavirus disease pandemic (Gold et al. 2023). Widespread antifungal resistance against commonly used drugs, particularly for *Candida auris*, worsens the current threat of fungal infection (Pristov and Ghannoum 2019; Miramon et al. 2023). For infections caused by Mucorales, drugs are generally insufficient, and radical surgical debridement is an important but disfiguring treatment.

The fungi that cause invasive lethal infections in humans are spread throughout the fungal kingdom, including members of the phyla Ascomycota (including *Candida albicans*, *C. auris*, *Aspergillus fumigatus, Histoplasma capsulatum, and Pneumocystis jirovecii*), Basidiomycota (*Cryptococcus neoformans*), and Mucoromycota (*Rhizopus arrhizus* a.k.a. *R. oryzae*). In previous studies, Trl1 homologs from the human ascomycete pathogens *C. albicans, A. fumigatus,* and *Coccidioides immitis* have been shown to complement *S. cerevisiae trl1Δ* strain (Remus et al. 2016), supporting that Trl1 function is conserved and a good candidate for being targeted by antifungal drugs.

In this study, we identified functional homologs of ScTrl1 in a broader range of human pathogenic fungi species that represent diverse evolutionary relatedness. The Trl1 homologs that have been shown to complement *trl1Δ* are all from other Ascomycete fungi. Here, we have identified ‘typical’ Trl1 orthologs in Ascomycota, Basidiomycota, and Mucoromycota pathogens that have all three heal and seal domains and are capable of complementing *trl1Δ*, supporting that Trl1 targeting drugs might be broad-spectrum antifungals. Interestingly, we identified an ‘atypical’ Trl1 ortholog in the order Mucorales (*R. arrhizus* and *Mucor circinelloides*). These Trl1 orthologs contain a Trl1 ligase domain, but lack CPD and kinase domains. These Mucorales Trl1 “sealing-only” orthologs can functionally complement defects in the ligase domain of ScTrl1 but not *trl1Δ*. Using orthologous mutation in the conserved nucleotide binding motif, we demonstrate that the sealing-only Trl1 of Mucorales is mechanistically similar to ScTrl1. Finally, the sealing-only Trl1 of Mucorales can perform the sealing step of *HAC1* mRNA splicing in yeast, indicating that both the known functions of Trl1 are retained. This suggests that drugs that target the ligase domain are likely to be broader-spectrum antifungals than those targeting the CPD or kinase domains. These findings imply that the Mucorales use kinase and CPD enzymes in a heal and seal pathway that are not orthologs to Trl1.

## Results

### Most pathogenic fungi contain a typical Trl1 ortholog, except the Mucorales

We sought to identify the orthologs of *S. cerevisiae* tRNA ligase (ScTrl1) in a diverse range of pathogenic fungi species, with a focus on pathogens that cause invasive disease with high morbidity and mortality. Two primary criteria were considered for identifying ScTrl1 orthologs: (a) the sequence-level similarities among the fungal *TRL1* genes with ScTRL1, and (b) the presence of three domains in the fungal *TRL1* genes. First, we compared the human pathogens that belong to the phylum Ascomycota. The Ascomycota are subdivided in three subphyla that each include important human pathogens. Among the subphylum Saccharomycotina, the genomes of two important human pathogens, *C. albicans* and *C. auris* (causative agents of candidiasis), encode ‘typical’ Trl1 orthologs with all three catalytic domains and an overall sequence similarity to ScTrl1 of 59% and 55%, respectively. Two representative fungi of the subphylum Pezizomycotina, *A. fumigatus* (causative agent of aspergillosis), and *H. capsulatum* (causative agent of histoplasmosis) also contain ScTrl1 orthologs with overall sequence similarity of 54% and 55%, respectively. Finally, the subphylum Taphrinomycotina includes one significant human pathogen, *P. jirovecii,* and its genome encodes a typical Trl1 ortholog with 52% similarity to ScTrl1. Thus, the genomes of all human pathogens within the ascomycetes encode a typical Trl1 ortholog. Within the phylum Basidiomycota, *C. neoformans* is the most important human pathogen and its genome encodes a ScTrl1 ortholog with 46% similarity to ScTrl1.

Strikingly, we could not identify a typical Trl1 ortholog in the genome of *Rhizopus* species, the most important pathogen within the Mucoromycota (e.g *R. arrhizus*) or in closely related species (e.g. *M. circinelloides*). Instead, the Trl1 orthologs in these species were much smaller and appeared to consist of only a ligase domain [Figure 1A]. These ligase domains were highly similar to that of ScTrl1 (55% and 49%). Similar sealing-only Trl1 homologs were encoded in other genomes of the order Mucorales, and all of them show sequence similarity with the ligase domain of ScTrl1. In contrast, in other Mucoromycota, we readily identified typical Trl1 orthologs that contain all three domains, (e.g. *Rhizophagus irregularis*, overall 52% similar to ScTrl1). This suggests that a typical Trl1 ortholog was present in the common ancestor of the Mucoromycota, Basidiomycota and Ascomycota, but was shortened after divergence of the Mucorales from the other fungi. Despite extensive searches, we were unable to identify any genes in the Mucorales that were orthologous to the CPD and kinase domains of Trl1.

**Figure 1:**
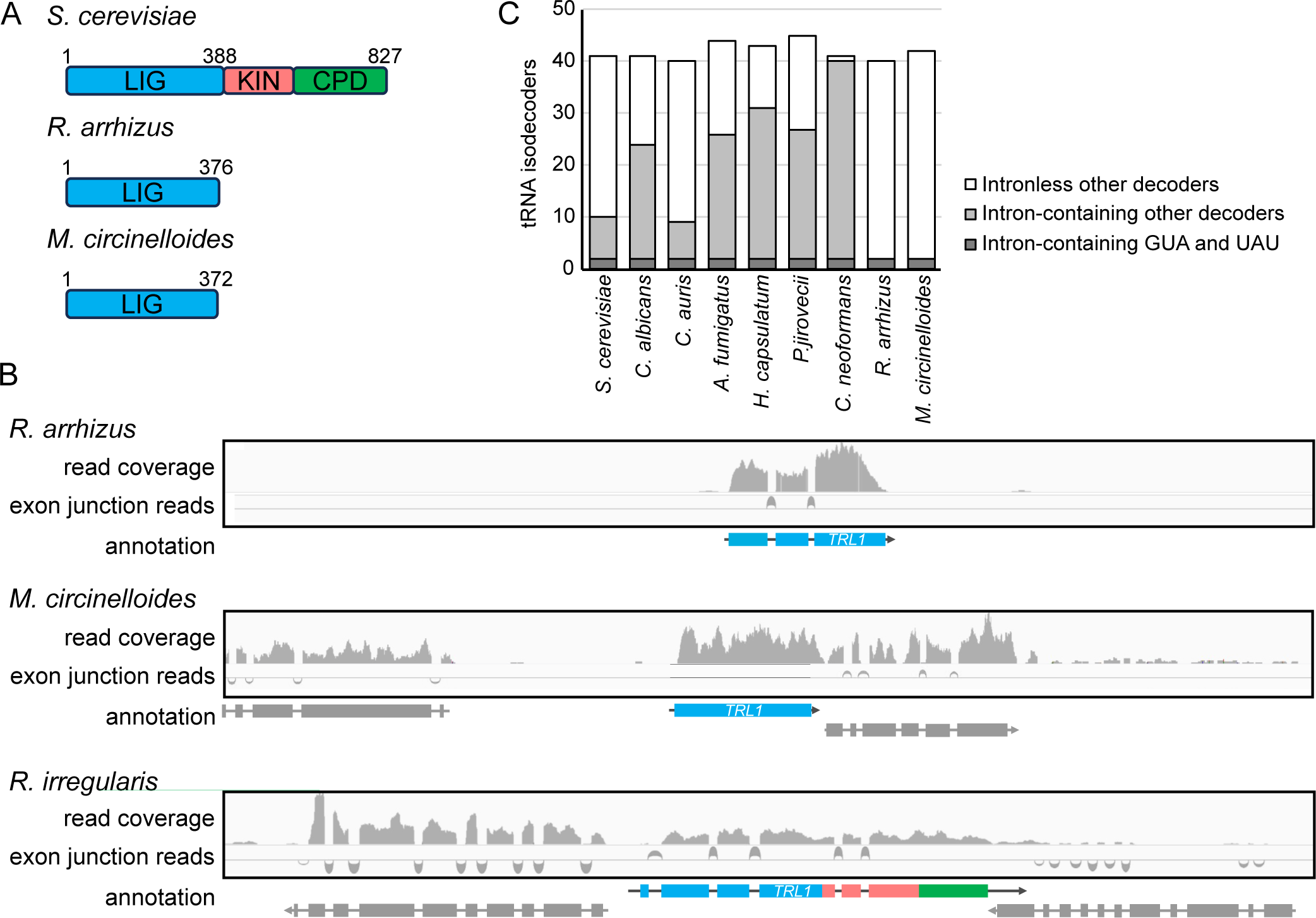
The Mucorales contain an atypical Trl1 ortholog with only the “sealing” domain. **(A)** Domain architecture of *S. cerevisiae* Trl1 and orthologs from two Mucorales (*R. arrhizus* and *M. circinelloides*). LIG; ligase domain, KIN; kinase domain, and CPD; cyclic phosphodiesterase domain. **(B)** The tRNA ligase of *M. circinelloides* and *R. arrhizus* express as sealing-only single domain Trl1s, whereas *R. irregularis* Trl1 contains three domains. Shown is RNA-seq read coverage of the *TRL1* orthologs of *M. circinelloides*, *R. arrhizus* (both Mucorales) and the related *R. irregularis* tRNA ligase. Colors reflect domains as in A. **(C)** The Mucorales contain a minimal set of intron-containing tRNAs. The number of intronless isodecoders (white), intron-containing isodecoders (grey), and intron-containing Tyr GUA (decoding UAC and UAU codons) and Ile UAU (decoding AUA codons) (dark grey) isodecoders in the genomes of major pathogenic fungi is shown.

To confirm that the Trl1 orthologs from *Rhizopus* and *Mucor* were truncated, we analyzed their gene structure by RNAseq. A re-analysis of publicly available RNAseq data confirms that transcripts of *R. arrhizus* and *M. circinelloides TRL1* orthologs encode only the ligase domain. In *M. circinelloides*, the ligase domain is encoded in a single exon that is not spliced with any other exon. In *R. arrhizus*, the ligase domain is encoded by three exons that are spliced together, but not to any other exons. In contrast, the four exons that encode the ligase domain of *R. irregularis* Trl1 are clearly spliced together with additional exons that encode the kinase and CPD domains [Figure 1B]. Overall our data suggest that most human fungal pathogens express a typical Trl1 ortholog, but the Mucorales instead express a ‘sealing-only Trl1’.

### The Mucorales contain a small number of tRNA introns and other components of a heal and seal tRNA splicing pathway

We investigated several possible explanations for the absence of a typical Trl1 from the Mucorales. First, we considered the possibility that a lack of tRNA introns in the Mucorales means they do not need a typical Trl1. Although all eukaryotes are thought to have tRNA introns, the proportion of tRNA genes that contain introns varies widely. We observed the same pattern when focusing on human fungal pathogens [Figure 1C] [Supplementary Table 1]. Eukaryotic genomes generally encode 42-50 distinct tRNAs (isodecoders) that each decode 1 to 4 different codons.

Each of these isodecoders can be encoded by a single tRNA gene or by multiple tRNA genes. If an isodecoder is encoded by multiple genes, typically either all of the genes for that isodecoder contain an intron, or none of them do. Among the fungal pathogens analyzed, we saw similar patterns, with *C. neoformans* containing the largest number of tRNA introns (92% of its tRNA genes). In contrast, *R. arrhizus* and *M. circinelloides* contain a relatively small number of intron-containing tRNAs (3% and 4% of its tRNA genes, respectively). Further analysis of isodecoder tRNAs in fungal pathogens revealed that all of the genes for tyrosine tRNA (which decodes both UAU and UAC) and for AUA isoleucine codons (but not AUU and AUC Ile codons) had an intron. Thus *R. arrhizus* and *M. circinelloides* require tRNA splicing to decode three of the 61 sense codons. While the genes for these same tRNAs contained introns in all fungal human pathogens, there was a wide variation in the number of other intron-containing tRNA isodecoders and genes [Figure 1C]. These observations rule out the possibility that the Mucorales do not need a typical Trl1 because they might lack tRNA introns.

The tRNA splicing endonuclease (TSEN) complex mediates the endonuclease cleavage of the intron-containing tRNA and comprises two catalytic subunits (Sen2 and Sen34) and two structural subunits (Sen15 and Sen54). The two catalytic subunits are more highly conserved at the sequence level, and we readily identified orthologs of both in the genome of all the major invasive fungal pathogens, including *R. arrhizus* and *M. circinelloides* [Supplementary Table 2]. The presence of both tRNA introns and a tRNA splicing endonuclease in the Mucorales strongly suggest that they must contain a fully functional tRNA splicing pathway.

One possible explanation for the absence of a three-domain Trl1 is that Mucorales acquired an RTCB-like ligase. We therefore searched their genomes but failed to detect any RTCB homologs. A third ligase (C12orf29) has recently been identified in humans and suggested to possibly function in tRNA splicing (Yuan et al. 2023), but we could not find any orthologs in the Mucorales genomes.

If the Mucorales indeed use the truncated Trl1 in a heal and seal tRNA splicing pathway, they should have homologs of Tpt1, the enzyme that removes the 2’ phosphate left after the sealing step. Indeed, the genomes of the Mucorales each include an obvious Tpt1 ortholog [Supplementary Table 2]. The most likely explanation of these bio-informatic analyses is that the Mucorales have functionally replaced the “healing” domains of Trl1 with an non-orthologous CPD and kinase. Similar replacements have previously been artificially achieved in *S. cerevisiae* (see discussion), but to the best of our knowledge has not been reported naturally.

### The sequence and predicted structure of sealing-only Trl1s of Mucorales suggest they are functional RNA ligases

The shorter tRNA ligase candidates of *R. arrhizus and M. circinelloides* contain 376 and 372 amino acids, respectively. Previous studies have identified that the N-terminal ligase domain of Trl1 orthologs contains six conserved peptide motifs (I, Ia, III, IIIa, IV, and V) that are found in the nucleotide-binding pocket (Wang and Shuman 2005). An amino acid sequence alignment reveals that these conserved motifs are present in *R. arrhizus* and *M. circinelloides* and therefore suggests that these proteins encode functional RNA ligases. [Supplementary Figure 1].

To gain insights into the structural features of Mucorales Trl1, we generated predicted structures of *R. arrhizus* and *M. circinelloides* Trl1 using both AlphaFold2 (Varadi et al. 2022) and Phyre tools (Kelley et al. 2015) [Figure 2 A-E]. Notably, although these tools use different approaches to predict structure, they give very similar results with high confidence. The predicted structures obtained from the analyses indicate that both Mucorales Trl1s share high structural similarities with the ligase domain of the available X-ray structure of the Ascomycete fungus *Chaetomium thermophilum* Trl1 (PDB: 6N0V, 6N0T, and 6N67) [Figure 2 A-C] (Banerjee et al. 2019; Peschek and Walter 2019). The secondary structures predicted by AlphaFold2 of the N-terminal subdomain of Mucorales Trl1 were superimposed with *C. thermophilum* Trl1 using the Matchmaker tool of Chimera [Figure 2 D-E]. The overall architecture and the catalytic pocket, including the conserved motifs, is conserved. The only notable difference in the N-terminal subdomains is a small N-terminal extension (∼30 amino acid residues) on the *C. thermophilum* protein that is distant from the catalytic pocket and unlikely to alter the function. The C-terminal subdomain of Mucorales Trl1 also shows high structural similarity to the *C. thermophilum* X-ray structure. In *C. thermophilum*, this domain consists of four alpha helices, and all four are conserved in the AlphaFold predicted Mucorales structures. The only notable difference between the AlphaFold2 predictions and the *C. thermophilum* structures is that AlphaFold2 predicts two short antiparallel beta strands inserted between the second and third alpha helix of the C-terminal subdomain. These strands are distant from the catalytic pocket and unlikely to alter the function of these proteins. Furthermore, AlphaFold2 predicts that the N- and C-terminal subdomains are oriented very similar to those in *C. thermophilum*.

**Figure 2:**
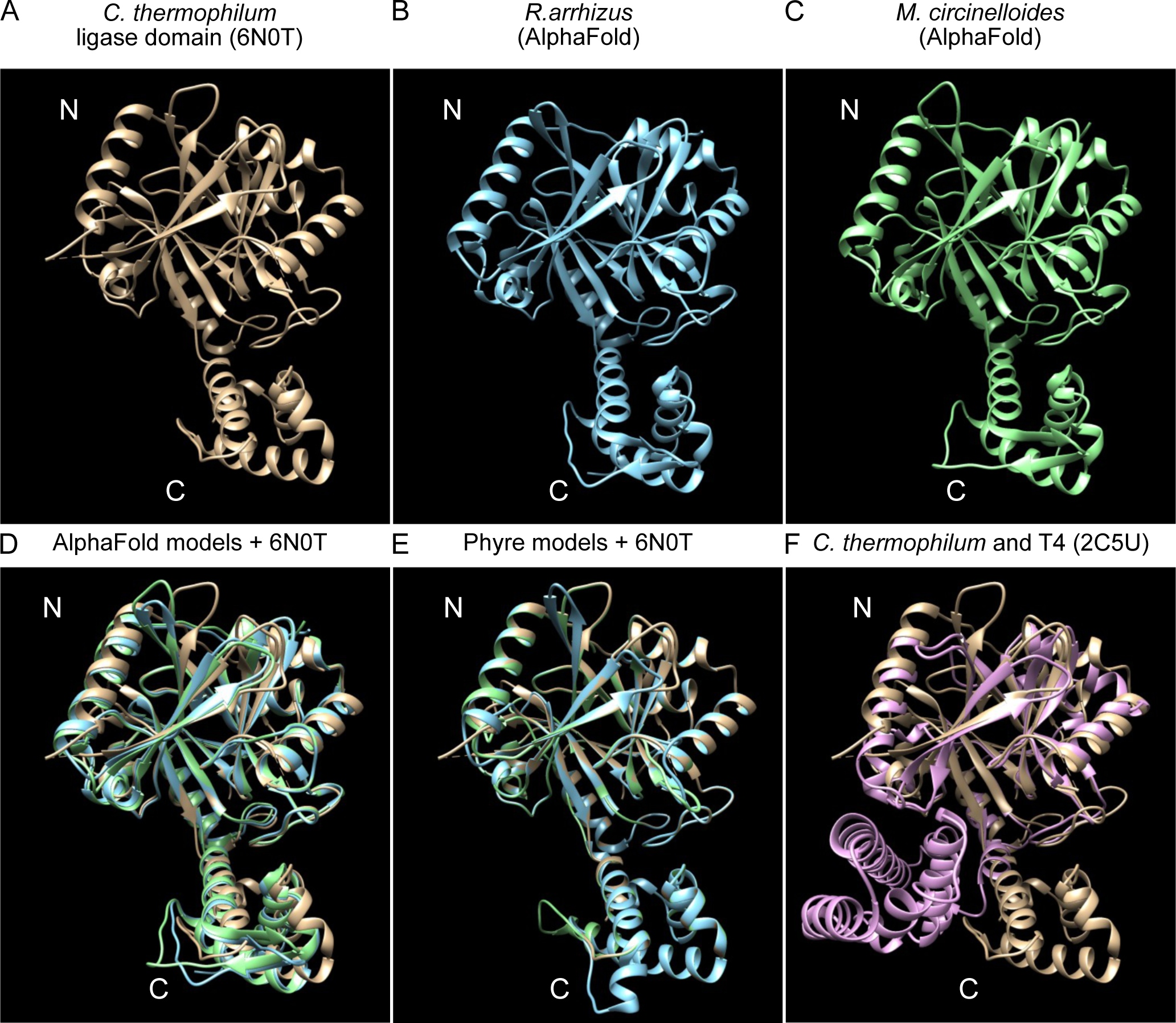
The predicted structures of Mucorales Trl1 resemble the ligase domain of fungal tRNA ligase. (**A)** X-ray crystal structure of the ligase domain of the ascomycete fungus *C. thermophilum* (6N0T). **B** The AlphaFold predicted structure of *R. arrhizus* sealing-only Trl1. (**B)** The AlphaFold predicted structure of *M. circinelloides* sealing-only Trl1. (**D)** Superimposition of X-ray crystal and AlphaFold predicted structures from A to C. Colors are as in A to C. For *R. arrhizus* RMSD is .913Å over 247 Cα atoms for M circinelloides it is .901Å over 244 Cα atoms (**E)** Superimposition of X-ray crystal and Phyre predicted structures. Colors are as in A to C. **F** Superimposition of X-ray crystal structures of Trl1 ligase from *C. thermophilum* (6N0T) and T4 RNA ligase (2C5U; plum colored). The N- and C-terminal subdomains are indicated in each panel.

To more objectively compare our AlphaFold2 predictions of *M. circinelloides* Trl1 and *R. arrhizus* Trl1 to known structures, we performed a DALI search of the protein data bank (PDB). For *M. circinelloides* and *R. arrhizus* Trl1, the top DALI recovered structure was the X-ray structure of the ligase domain of the *C. thermophilum* (PDB 6N0V) with a z-score of 41.8 and 41.3, respectively (Banerjee et al. 2019). The second most related structure from the DALI search was the X-ray structure of T4 RNA Ligase (Rnl1) (PDB 2C5U) (El Omari et al. 2006). A key distinction between Trl1 and T4 Rnl1 is that Trl1 uses a substrate with a 2’PO_4_ 3’OH while T4 Rnl1 uses a substrate with 2’OH 3’OH (Schwer et al. 2004; Banerjee et al. 2019). This biochemical difference is mediated by a C-terminal extension on the Trl1 ligase domain (Banerjee et al. 2019). The 6N0T structure identifies key residues that recognize the 2’PO4 (N150, H227, R334 and R337), and each of these residues is conserved in the *Rhizopus* and *Mucor* proteins and positioned in the same place in the AlphaFold2 models. The R334 and R337 residues are part of a YRxxR motif in the C-terminal subdomain, and the T4 Rnl1 C-terminal subdomain does not contain a similar motif. The T4 Rnl1 subdomain is also positioned differently relative to the N-terminal subdomain [Figure 2F]. Taken together, the predicted structures of Mucorales Trl1 suggest these proteins are active RNA ligases that require a 2’PO_4_.

### The sealing-only Trl1s of Mucorales functionally complement the ScTrl1 ligase domain

In order to determine whether the Trl1 orthologs from invasive pathogenic fungi are functional, we expressed Trl1 orthologs in a *S. cerevisiae trl1Δ* strain. A set of plasmid shuffle assays allowed for the selection of the yeast cells that express fungal Trl1 orthologs and have lost the wild-type ScTrl1 plasmids [Figure 3A]. Each of the tested three-domain containing ‘typical fungal Trl1s’ from pathogenic fungi complemented *trl1Δ*. Specifically, replacing Trl1 with the Trl1 orthologs from the closely related *C. albicans* and *C. auris* resulted in growth that resembled wild type. Complementation with the ortholog of the more distantly related Ascomycota *A. fumigatus* or *H. capsulatum* also resulted in growth, although at a reduced rate. We were unable to include *P. jirovecii* in this experiment because this gene proved toxic during cloning in *E. coli.* This complementation by orthologs from other Ascomycota confirms and extends previous reports (Remus et al. 2016). Remarkably, the full-length Trl1 orthologs from the Basidiomycota *C. neoformans* and the Mucoromycota *R. irregularis* can also complement *trl1Δ* and resulted in near-normal growth rate. This indicates that all of the typical three-domain fungal Trl1s can perform tRNA ligation in budding yeast. In contrast, as expected the truncated Trl1 orthologs from *R. arrhizus* and *M. circinelloides* failed to complement *trl1Δ* and behaved similar to an empty vector control.

**Figure 3:**
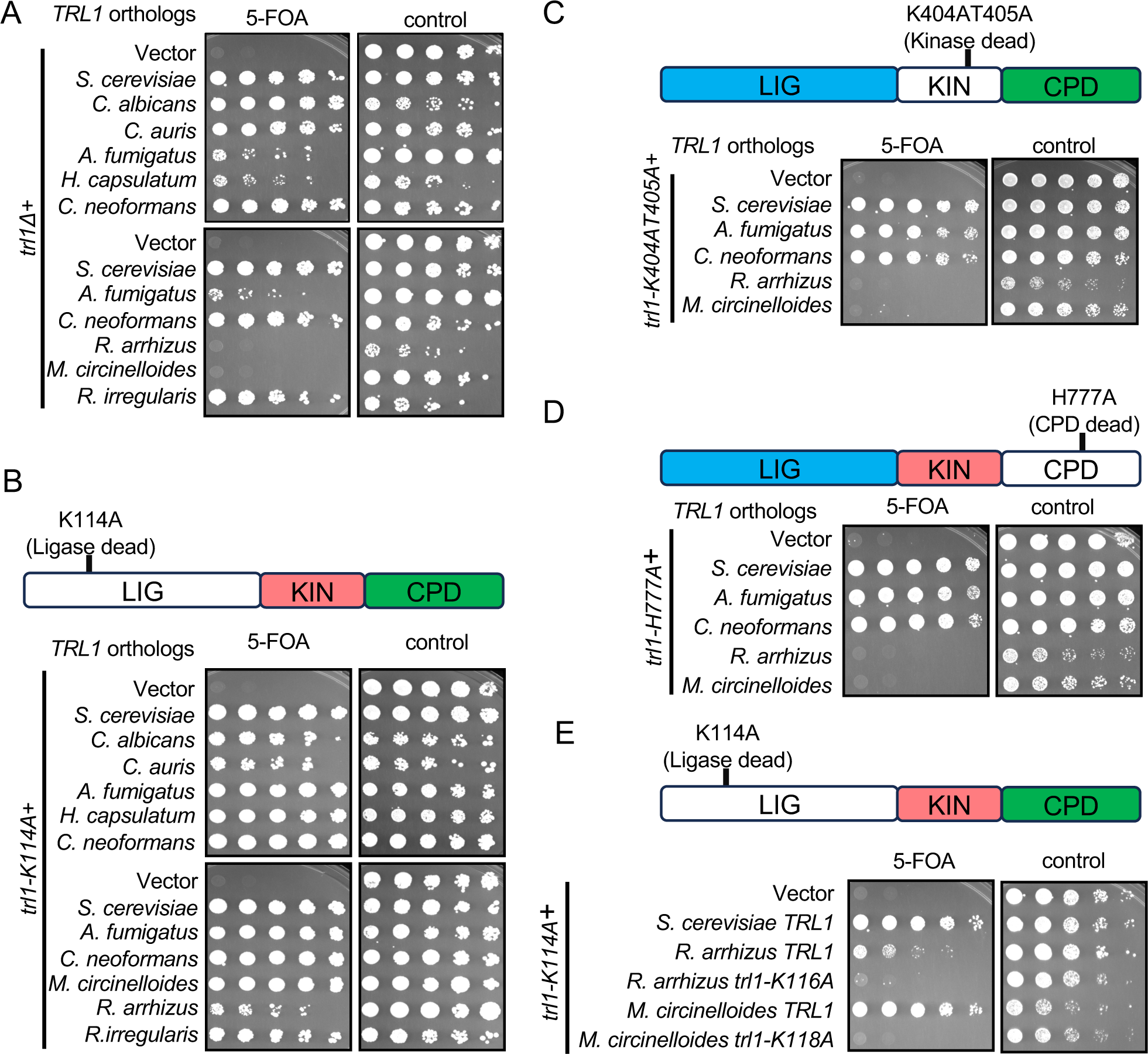
Complementation of Trl1 defects by orthologs from fungal pathogens. **(A)** The Trl1 orthologs of representative pathogenic fungi complement *trl1Δ*, except the sealing-only Trl1s of Mucorales. The indicated Trl1 orthologs were expressed in *S. cerevisiae trl1Δ* [*TRL1*, *URA3*] strain background and subjected to serial dilution growth assay on media containing 5-FOA (selecting against the *TRL1* plasmid) or control media. (**B-D)** The Mucorales Trl1 can complement a defect in the ligase (LIG) domain of ScTrl1 (B) but is unable to complement the defects in the kinase (KIN; C) or cyclic phosphodiesterase (CPD; D) domain of ScTrl1. The indicated Trl1 orthologs were expressed in a strain that contains a *URA3* plasmid with a wild-type *TRL1* gene and another plasmid with either the sealing-defective *trl1-K114A* (B), the kinase-defective *trl1-K404AT405A* (C), or the cyclic phosphodiesterase-defective *trl1-H777A* (D) subjected to serial dilution growth assay on media containing 5-FOA (selecting against the *TRL1* plasmid) or control media. (**E)** A conserved lysine in the active site is critical in the Mucorales Trl1s. Amino acid changes orthologous to scTrl1-K114A were introduced in Mucorales Trl1 and subjected to plasmid shuffle assay in the sealing-defective ScTrl1 strain as in B.

We postulated that because the identified Mucorales Trl1s only contain the ligase domain, they could only complement the exon ligation step (sealing step) but not the healing steps of the fungal type tRNA splicing pathway. A conserved lysine residue in position 114 of ScTrl1 is required for the function of the ligase domain. Amino acid change of the K114 residue to alanine (K114A) exhibits defects in the sealing function of Trl1 and is lethal (Sawaya et al. 2003). We therefore performed a plasmid shuffle assay to test whether Trl1 orthologs of pathogenic fungi can functionally complement a ligase domain defect (*trl1-K114A*). As expected, all of the fungal Trl1 orthologs that complement *trl1Δ* also complemented *trl1-K114A*. Importantly, the sealing-only Trl1s from Mucorales can complement this ligase-dead ScTrl1 [Figure 3B]. As expected, we confirmed that the Mucorales Trl1s cannot perform the ‘healing’ reactions of tRNA splicing pathway in budding yeast: they do not complement mutations in either the ScTrl1 kinase domain (*trl1-K404AT405A*) or the CPD domain (*trl1-H777A*) [Figure 3C-D].

We examined whether the Mucorales Trl1s use the same catalytic lysine as ScTrl1. Sequence alignment identified the corresponding conserved lysine residues as K116 for *R. arrhizus* and K118 for *M. circinelloides*. A plasmid shuffle experiment indicates that changing these lysine residues to alanine in either of the Mucorales proteins disrupted the ability to complement the defect in the ScTrl1 ligase domain [Figure 3E]. Overall, these data indicate that Mucorales Trl1s are functional tRNA ligases and are able to perform the tRNA sealing step but not the tRNA healing steps.

### Mucorales Trl1 can perform *HAC1* mRNA splicing during the Unfolded Protein Response

As discussed in the introduction, in addition to tRNA splicing, Trl1 takes part in the non-conventional *HAC1* mRNA splicing mechanism during the Unfolded Protein Response (UPR) that is initiated by Ire1. The UPR plays a critical role in the stress tolerance and virulence of pathogenic fungi (Richie et al. 2009; Cheon et al. 2011; Askew 2014; Krishnan and Askew 2014). During infection, fungal pathogens encounter several biotic and abiotic stresses in the host. Thus, an increased protein folding capacity in the endoplasmic reticulum (ER) via UPR signaling is required to overcome the host stresses associated with the increased pathogenicity of the fungal pathogens.

The Mucorales genomes contain an obvious ortholog for Ire1, but Hac1 orthologs are poorly conserved in sequence and difficult to detect across large evolutionary distance. Nevertheless, we identified a likely Hac1 ortholog in *R. arrhizus* (hypothetical protein G6F37_009984;) and RNAseq analysis identified an unannotated non-canonical intron [Figure 4A].

**Figure 4:**
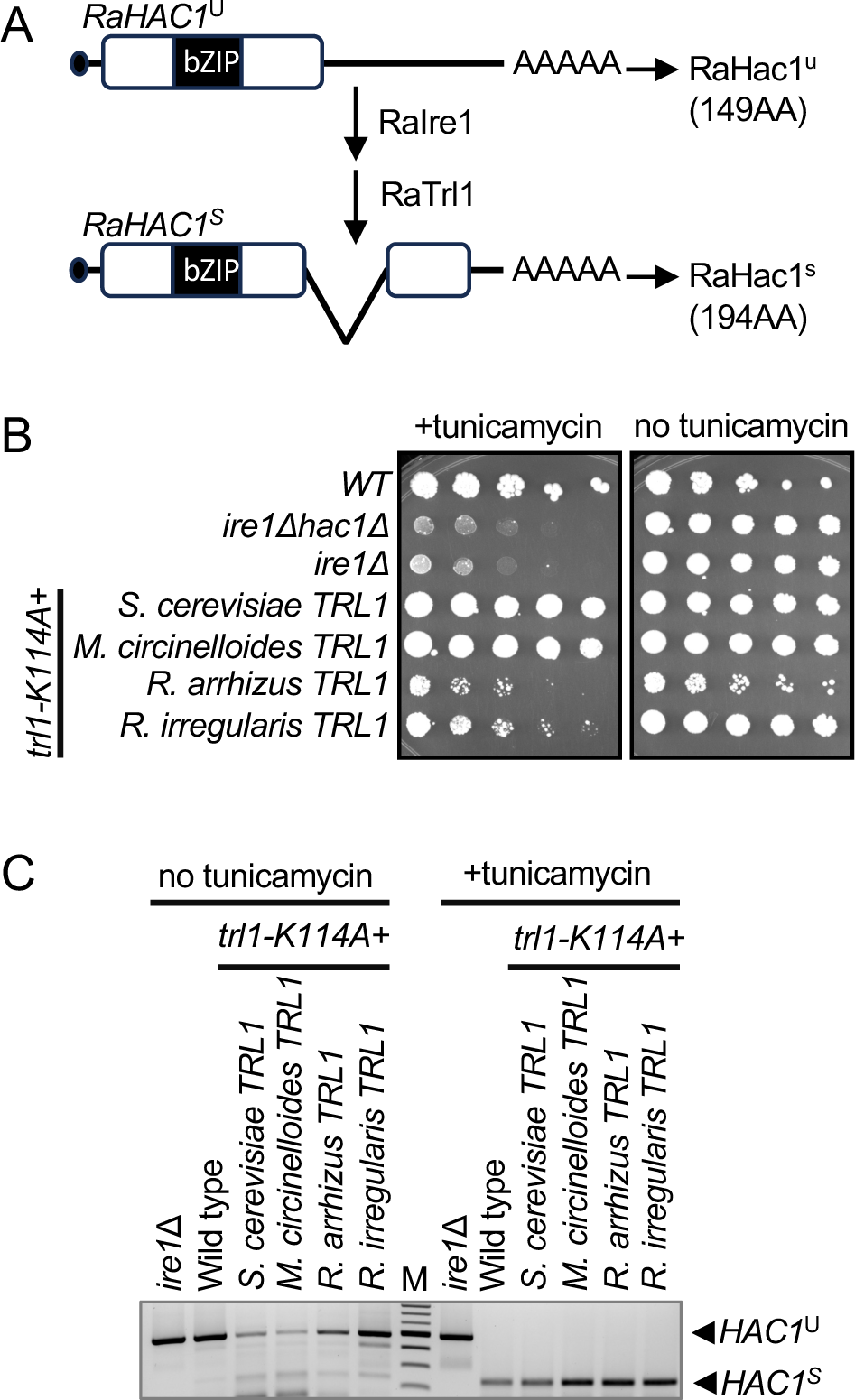
Mucorales sealing-only Trl1s can perform *HAC1* splicing during the Unfolded Protein Response (UPR). **(A)** *R. arrhizus* appears to have an Ire-dependent intron in *HAC1* mRNA (encoding the 149 amino acid hypothetical protein G6F37_009984). Sequence similarity to Hac1 is highest in the indicated bZIP domain. Depicted is the unspliced (RaHAC1^u^) and spliced (RaHAC1^s^) mRNA structure based on aligning RNAseq data (SRR1013749) to the genome with RNA STAR, and the predicted function of Ire1 and sealing-only Trl1 in generating spliced *HAC1* mRNA. (**B)** The *M. circinelloides*, *R. arrhizus,* and *R. irregularis* Trl1 were expressed in *S. cerevisiae trl1Δ* [*trl1-K114A*] [*TRL1*, *URA3*] strain background and subjected to plasmid shuffle assay in the presence or absence of 0.2 µg/mL tunicamycin. The wild-type (WT), *ire1Δhac1Δ,* and *ire1Δ* strains were used as a control. (**C)** RT-PCR of *HAC1* indicates that upon UPR induction, the Mucorales Trl1 gives rise to spliced *HAC1* mRNA (*HAC1^S^*). The *HAC1* splicing defective *ire1Δ* strain was used as a control, which only produced the unspliced *HAC1* mRNA (*HAC1^U^*).

To test whether *R. arrhizus* and *M. circinelloides* Trl1 are able to perform *HAC1* mRNA splicing in budding yeast, we repeated the plasmid shuffle assay in the sealing-defective ScTrl1 strain (*trl1-K114A*) without or with 0.2µg/mL tunicamycin. Tunicamycin causes unfolded proteins to accumulate in the ER; thus, the UPR is required for viability in the presence of 0.2µg/mL tunicamycin. Indeed, *ire1Δ* and *ire1Δhac1Δ* control strains failed to grow in the presence of tunicamycin. Importantly, the strains that express Trl1 from the Mucorales and a sealing-dead ScTrl1 grew in the presence of tunicamycin and thus complemented the UPR defect [Figure 4B], although the *M. circinelloides* ortholog performed better than the *R. arrhizus* ortholog. As a control, the three-domain Trl1 from the Mucoromycota *R. irregularis* also complemented the UPR defect.

To confirm that growth on tunicamycin reflected the ability to splice *HAC1*, we assayed *HAC1* mRNA splicing directly by RT-PCR with primers that span the intron [Figure 4C]. When the cells were grown without UPR stress, the major RT-PCR product was the size expected for unspliced *HAC1* mRNA. RT-PCR of the strains that express Mucoromycota Trl1s in addition to the ScTrl1 sealing-dead mutant resulted in a major product of 247 bp after UPR induction, reflecting removal of the *HAC1* intron. The UPR defective *ire1Δ* strain used as a control produced only unspliced *HAC1* mRNA. Taken together, these results indicate that *R. arrhizus* and *M. circinelloides* Trl1s are able to carry out the sealing step of *HAC1* splicing during UPR stress when expressed in *S. cerevisiae*.

## Discussion

In this study we show that the tRNA ligase (Trl1) orthologs from a diverse range of clinically relevant fungal pathogens are functionally conserved and can replace the Trl1 enzyme in *S. cerevisiae*. These fungal species contain a subset of tRNA genes that require splicing to produce functional tRNA molecules. Specifically, in each species the tRNA-Tyr-GUA and tRNA-Ile-UAU tRNA genes contain an intron, while additional tRNA genes contain an intron in a subset of pathogenic fungi. This indicates that tRNA splicing is essential in all invasive fungal human pathogens. Alongside Trl1 orthologs, the presence of Tpt1 orthologs and the absence of alternative ligases (RTCB or C12orf29 homologs) confirm that the ‘healing and sealing’ mechanism is the prevalent mode of tRNA exon ligation in fungi. There is an urgent need for novel antifungal drugs. There are very few enzymes that are essential across the fungal kingdom, but absent in humans. The heal and seal pathway is a prominent exception and thus an attractive target for antifungal drugs.

Pathogenic fungi including *A. fumigatus* and *C. neoformans* utilize the UPR to fine-tune protein folding capacity in the ER, which is linked to virulence, membrane homeostasis, cell wall homeostasis and anti-fungal drug resistance (Richie et al. 2009; Cheon et al. 2011; Askew 2014; Krishnan and Askew 2014; Weichert et al. 2020). Thus, targeting the fungal UPR has been suggested as a potential alternative strategy for antifungal development. The advantage of Trl1 as a drug target is that a Trl1 inhibitor would inhibit both tRNA splicing and the UPR in pathogenic fungi.

Unlike other representative fungi, Mucorales (*e.g. R. arrhizus* and *M. circinelloides*) appear to have separate sealing and healing enzymes to carry out tRNA exon ligation. Our identified Trl1 orthologs in Mucorales are functional ligases that can carry out the sealing reactions in both known Trl1 functions: tRNA and *HAC1* mRNA splicing. However, the lack of the Trl1 kinase and CPD domains in these orthologs suggests that they likely act with some other polynucleotide kinase and phosphodiesterase that carry out the healing steps of tRNA and *HAC1* splicing. There is precedence of this both in nature and in previous experiments. Englert et al. (2010) described two Trl1 orthologs in the chordate *Branchiostoma floridae* (Florida lancelet) (Englert et al. 2010) that are each truncated. One of the proteins contains only the sealing domains, while the other contains the healing domains. The *B. floridae* genome also encodes RTCB and C12orf29 orthologs (XP_035676627 and XP_035678444). Which of these proteins splice the 42 annotated tRNA introns in *B. floridae* (http://gtrnadb.ucsc.edu/) remains unknown. Despite extensive efforts, we failed to detect proteins in the Mucorales that are orthologous to the healing domains of Trl1, making the Mucorales distinct from *B. floridae*. Experimentally, the healing domains of Trl1 in yeast can be replaced by the distantly related mammalian RNA 5’-kinase CLP1 and 2’,3’ cyclic nucleotide phosphodiesterase (CNP) (Ramirez et al. 2008; Schwer et al. 2008). We propose that the Mucorales similarly use distantly related healing enzymes in their healing and sealing pathway, but the identity of these enzymes remains unknown. Because the Mucorales proteins can seal the 2’ PO_4_ and 5’ PO_4_ ends generated by yeast healing domains, the unknown kinase and CPD likely also generate 2’ PO_4_ and 5’ PO_4_ ends.

Our study supports the possibility of targeting the tRNA exon ligation mechanism for antifungal development. The ligase domain of fungal Trl1 is conserved in all of the important invasive human pathogens, including the Mucorales. Thus, a drug that inhibits the function of the Trl1 ligase domain should be a broad-spectrum antifungal against these fungal pathogens. In contrast, drugs that target the kinase and CPD domains are less likely to be effective against the Mucorales. However, an antifungal that targets Ascomycete and Basidiomycete Trl1s would still be useful, as these taxa include the most urgent fungal threats (https://www.cdc.gov/drugresistance/biggest-threats.html) and account for most of the cost of fungal infections (Benedict et al. 2019).

## Materials and methods

### Identification of fungal Trl1 genes

To identify Trl1 candidates in representative pathogenic fungi, extensive BLAST and PSI BLAST searches were performed starting with either ScTrl1 or the *R. irregularis* ortholog. The *TRL1* genes were identified based on the similarity of the domain composition and the sequence. Orthologs of Sen2, Sen34, and Tpt1 were similarly identified with BLAST and PSI BLAST. The search for alternative ligases used human RTCB, human C12ORF29, and *E. coli* RtcB protein sequences.

To identify the correct exon junctions, publicly available RNA sequencing reads from *R. arrhizus* strain 99-892 (SRR1013749 and SRR1013747), *M. circinelloides* strain R7B (SRR1611143, SRR1611144, and SRR1611152), and *R. irregularis* (SRR14294959 and SRR14294960; (Dallaire et al. 2021)) were aligned to the corresponding reference genome assemblies (GCA_024220505.1, GCA_001638945.1, and GCF_000439145.1 (Tisserant et al. 2013), respectively) using RNA STAR (Dobin et al. 2013).

tRNA annotations and their introns are from the genomic tRNA database (http://gtrnadb.ucsc.edu/index.html); for species not included, they were identified using tRNAscan-SE 2.0, the same algorithm used for the genomic tRNA database (Chan and Lowe 2016; Chan et al. 2021).

### Plasmids

Plasmid used are listed in Supplemental table 3. Each of the *TRL1* orthologs was cloned into p425GPD (Mumberg et al. 1995) using Gibson assembly. For *C. albicans* and *C. auris*, the single exon genes were PCR amplified from genomic DNA (a kind gift from Michael Lorenz). For the other species, we did not have ready access to genomic DNA or cDNA and therefore the *TRL1* ORF was codon optimized for expression in *S. cerevisiae* and synthesized by GenScript.

The catalytic mutants for McirTRL1 (Mcir trl1-K118A) and RarrTRL1 (Rarr trl1-K116A) plasmids were created using the QuikChange Lightning Site-Directed Mutagenesis Kit following manufacturer’s instructions.

The catalytic mutants for ScTRL1 (trl1-K114A, trl1-H777A and trl1-K404A-T405A) were generated by Gibson assembly into pRS413 (Sikorski and Hieter 1989; Sawaya et al. 2003).

All plasmids were sequence verified either by sequencing the whole plasmid using nanopore sequencing (Plasmidsaurus) or by sequencing the insert using Sanger sequencing (Genewiz).

### Complementation of fungal Trl1s in *trl1Δ* yeast

The yeast strains used are listed in supplemental table 4. To identify whether the Trl1 of representative fungi can complement the loss of ScTrl1, the corresponding *TRL1* expression plasmids of fungi were transformed into S. *cerevisiae trl1Δ* [*TRL1, URA3*] strain. This parent strain was created from the S. *cerevisiae* heterozygous knockout library by transforming *TRL1, URA3* plasmid (pAV1511) and sporulating the transformants. The resulting strain was *matA*, *ura3-Δ0, leu2-Δ0, his3-Δ1, lys2-Δ0, trl1Δ::NEO, [TRL1,URA3]* and unable to grow on 5-FOA-containing media.

The transformants for yeast cells containing *TRL1* expression plasmids were selected in the SC media lacking leucine and uracil (SC-Leu-Ura, Sunrise Science). A plasmid shuffle experiment was performed, where the serially diluted yeast strains were spotted on 5-FOA-containing media that selects for yeast cells that have lost the wild-type *ScTRL1, URA3* plasmid and contains only the *TRL1* of corresponding fungi (Boeke et al. 1987). Growth was recorded after 4 days at 30^°^C.

To identify the ability of the fungal Trl1 to complement the defects in individual domain (LIG, KIN, CPD) function of ScTrl1, the corresponding *TRL1* plasmids were transformed into *trl1Δ* strains that contain sealing-dead, CPD-dead or kinase-dead Trl1s in pRS413 (CEN, *HIS3*) plasmids in addition to the *TRL1* gene on a CEN, *URA3* plasmid. The transformants were selected on SC-Leu-His media (Sunrise Science), and a plasmid shuffle assay was performed as described above.

To identify the ability of the fungal Trl1 to complement the Trl1 function in the unfolded protein response, *trl1Δ* yeast cells containing a *LEU2* plasmid expressing the *TRL1* of other fungi, a *HIS3* plasmid expressing *Sc trl1-K114A* and a *URA3* plasmid expressing wild-type *ScTRL1* were serially diluted and spotted on 5-FOA-containing media supplemented with or without 0.2 µg/mL tunicamycin (Sigma-Aldrich). Growth was recorded after 4 days at 30^°^C. The control *ire1Δ* strain was obtained from the knock out collection(Giaever et al. 2002). The control *ire1Δhac1Δ* strain (YAv4415) was generated by standard genetic crosses of single mutants obtained from the knock out collection. The *hac1Δ::KANMX* allele was converted to *hac1Δ::NATMX* using plasmid p4339 (obtained from Charles Boone).

### RNA extraction and RT-PCR

Overnight cultures of yeast strains were diluted to ∼0.1 OD_600_ in YPD and grown to ∼0.6 OD_600_ at 30^°^C. UPR induction was carried out by adding 2.5 µg/mL tunicamycin for 2 hours. The cells were harvested, and total RNA was extracted using the hot phenol method (He et al. 2008). The RNA samples were treated with DNase I (Invitrogen), and cDNA was synthesized with SuperScript™ II Reverse Transcriptase (Invitrogen). The *HAC1* spliced and unspliced fragments were amplified with intron-spanning primers (ACCTGCCGTAGACAACAACAAT and AAAACCCACCAACAGCGATAAT) and analyzed on a 2% agarose gel (Cherry et al. 2019).

## Acknowledgements

This work was funded by National Institutes of Health (NIH) grant R35GM141710 to A.v.H. and a Dr. John J. Kopchick Research grant from the UT MD Anderson Cancer Center UTHealth Houston Graduate School of Biomedical Sciences to K.S.A. We thank Sean Johnson for help with AlphaFold2 predictions and Michelle Steiger, Mike Lorenz, Sean Johnson, and members of the van Hoof lab for insightful comments.

## Notes

### Competing Interest Statement

The authors have declared no competing interest.

